# Herbarium specimens reveal links between *Capsella bursa-pastoris* leaf shape and climate

**DOI:** 10.1101/2024.02.13.580180

**Authors:** Asia T Hightower, Daniel H Chitwood, Emily B Josephs

## Abstract

- Studies into the evolution and development of leaf shape have connected variation in plant form, function, and fitness. For species with consistent leaf margin features, patterns in leaf architecture are related to both biotic and abiotic factors. However, for species with inconsistent leaf margin features, quantifying leaf shape variation and the effects of environmental factors on leaf shape has proven challenging.
- To investigate leaf shape variation in species with inconsistent shapes, we analyzed approximately 500 digitized *Capsella bursa-pastoris* specimens collected throughout the continental U.S. over a 100-year period with geometric morphometric modeling and deterministic techniques. We generated a morphospace of *C. bursa-pastoris* leaf shapes and modeled leaf shape as a function of environment and time.
- Our results suggest *C. bursa-pastoris* leaf shape variation is strongly associated with temperature over the *C. bursa-pastoris* growing season, with lobing decreasing as temperature increases. While we expected to see changes in variation over time, our results show that level of leaf shape variation is consistent over the 100-year period.
- Our findings showed that species with inconsistent leaf shape variation can be quantified using geometric morphometric modeling techniques and that temperature is the main environmental factor influencing leaf shape variation.

## Introduction

It is crucial to understand how complex traits relate to environmental variation, especially in the context of a rapidly changing climate (Anderegg, 2015; Cochrane *et al*., 2015; Henn *et al*., 2018; Moran *et al*., 2016)). Leaf shape is a complex trait with variation at developmental, environmental, and phylogenetic levels (Chitwood *et al*., 2014a,b; Chitwood & Sinha, 2016; Lin *et al*., 2020). For decades, the molecular and morphometric study of leaf shape and its effects on leaf function and plant fitness (Winn, 1999) have been important for advancing crop breeding (Andres *et al*., 2016; Hao *et al*., 2022), reducing pesticide use (de la Paz Pollicelli *et al*., 2018; Rivero-Lynch *et al*., 1997), and ultimately improving human health (Broadley & White, 2010; Key *et al*., 2008). Numerous paleoclimatic and common garden studies have shown that the size and shape of leaves often correlates with temperature and soil moisture on both the local and global scales (Dolph & Dilcher, 1980; Gregory-Wodzicki, 2000; Huff *et al*., 2003; Feild *et al*., 2005; Gleason *et al*., 2018; Royer *et al*., 2008)). In addition, leaf shape variation is often associated with fitness variation (Bright & Rausher, 2008; Ferris, 2019; Richards *et al*., 2019).

Leaf shape is a complex trait that is affected by genetic and environmental factors (Chitwood & Sinha, 2016). Leaf shape is frequently defined by its leaf margin dissections (lobing) (Peppe *et al*., 2011). Lobed leaves are simple leaves with leaf margin dissections, making them distinct from compound leaves, which have multiple subunits (’leaflets’) and discontinuous lamina (Runions *et al*., 2017; Bar & Ori, 2014). Lobe characteristics are often related to abiotic factors. Generally, increased lobing promotes photosynthesis (Baker & Myhre, 1969; Bhagsari & Brown, 1986; Smith *et al*., 1997; Kern *et al*., 2004; Nicotra *et al*., 2008; Tsukaya, 2018), water transportation (Passioura, 1988; Zwieniecki *et al*., 2004; Katifori, 2018; Ding *et al*., 2020; Sakurai & Miklavcic, 2021), and gas exchange (Araus *et al*., 1986; Pettigrew *et al*., 1993; Bednarz & van Iersel, 2001; de Boer *et al*., 2016; Harrison *et al*., 2020; Tamang *et al*., 2023). Overall, in warm environments, leaves are typically less lobed than leaves in cool environments (Dolph & Dilcher, 1980; Gregory-Wodzicki, 2000; Royer *et al*., 2008).

Many plant species have regular leaf shapes. For example, grape vine (*Vitis vinifera*) leaves are palmate and include five major veins (Chitwood *et al*., 2014b), *Arabidopsis thaliana* leaves are simple with unbroken leaf margins or serrations (Runions *et al*., 2017; Barkoulas *et al*., 2008)) and Cotton (*Gossypium hirsutum L.*) leaves include four major shapes that show differences in carbon fixation depending on other environmental conditions (de Boer *et al*., 2016; Andres *et al*., 2017; Pettigrew & Gerik, 2007). However, many species do not have consistent leaf shapes, especially in varying environments and we know significantly less about the development and evolution of leaf shape in species with inconsistent lobing (Kusi & Karsai, 2020; Geeta *et al*., 2012). In addition, it is more challenging to study shape in plants with inconsistent lobing: the lack of consistent and/or homologous points on leaves that have variable lobe numbers, lobe depths, and lobe angles makes comparisons among shapes difficult (Valenzuela *et al*., 2011; Chitwood & Otoni, 2017). Therefore, it is important that we can reliably investigate how leaf shape varies among species with inconsistent lobing across both evolutionary and ecological gradients (Bensmihen *et al*., 2008). As rising temperatures and increased *CO*_2_ become more prevalent (Pritchard *et al*., 1999; Royer, 2012), understanding how species with inconsistent lobing are affected by and can be adapted to combat these environmental changes becomes increasingly important.

Geometric morphometrics is an increasingly popular technique used to summarize shape in terms of a multidimensional landmark configuration, where shapes exist as Cartesian coordinates that can be transformed and compared across two and three dimensions (IIa & Mikeshina, 2002; Adams *et al*., 2004; Mitteroecker & Gunz, 2009; Webster & Sheets, 2010; Polly & Motz, 2016). For many species, the lack of consistency in trait features such as leaf margin lobing or serrations presents challenges in comparing landmarks within species and between species, as these homologous points may not exist. We address this issue with pseudo-landmarks: points placed between landmarks to estimate curves and to create more continuous representations of shape (Parsons *et al*., 2009; Budd, 2021).

Herbaria, or plant collections, are key resources of trait variation for a wide range and diversity of species over both time and geographic space (Moeller *et al*., 2007; Moloney *et al*., 2009; Menne *et al*., 2012; Gutaker *et al*., 2017; Chen *et al*., 2018; Borges *et al*., 2020; de Villemereuil *et al*., 2016; Sang-Hun, 2022). Herbarium collections span the U.S. civil war era to post-pandemic America (James *et al*., 2018; Lavoie, 2013; Park *et al*., 2023). Specimens in herbarium collections, which can include whole pressed plants, seeds, fruits, and much more, are a snapshot of the world at the time of collection (de Villemereuil *et al*., 2022; Willis *et al*., 2017; Heberling *et al*., 2019; James *et al*., 2018). A major strength of herbarium specimens is that they provide a view of plant traits from their natural environment, allowing researchers to assess trait changes in time and space (Willis *et al*., 2017; Lang *et al*., 2019). Through the use of genomic, digitization, and bioinformatics techniques, research with herbarium specimens has increased exponentially (Davis, 2023; Besnard *et al*., 2018; Miller-Rushing *et al*., 2004). Recent work using herbarium specimens has shown that comparisons of the association between traits and the climate across all years, some years, and the climate in the specific year of collection can be used to disentangle genetic and plastic trait changes (Wu & Colautti, 2022; Lang *et al*., 2019). Here we use herbarium leaf shape data to measure and compare leaf shape variation in *Capsella bursa-pastoris*, a species with well documented high variation in leaf shape and highly inconsistent leaf margin architecture.

*Capsella bursa-pastoris* a weedy allotetraploid in the Brassicaceae family, is a model system for investigating within-species leaf shape variation across a large environmental range (Aksoy *et al*., 1999; Shull, 1909). *C. bursa-pastoris* is found in most regions of the world (Choi *et al*., 2019; Cornille *et al*., 2022; Neuffer *et al*., 2018; Wesse *et al*., 2021) and has incredible variation in leaf shape (Neuffer, 1990; Hurka & Neuffer, 1997; Shull, 1909; Iannetta *et al*., 2007). Traditionally, phenotyping of leaf shape for *C. bursa-pastoris* leaves has used plant material collected from common garden environments and dichotomous leaf keys as an identification tool. These common garden studies have found that *C. bursa-pastoris* leaves can be categorized into shapes, referred to here as the ‘Shull types’ (Shull, 1909) or ‘Ianetta types’ (Iannetta *et al*., 2007) and suggested that there is a Mendelian genetic basis for leaf shape distribution following a temperature and elevation gradient (Aksoy *et al*., 1999; Neuffer, 1990). However, many studies find leaves that do not fit into one of the four Shull types (Aksoy *et al*., 1999; Shull, 1909; Begg *et al*., 2012). In addition, information from common garden experiments alone may miss key morphological information (Moloney *et al*., 2009; de Villemereuil *et al*., 2022, 2016) and assigning leaf shapes with dichotomous keys depends on the researcher’s judgment and therefore can be a subjective determination (Wiemann *et al*., 1998; Thyagharajan & Kiruba Raji, 2019; Li *et al*., 2020). Instead, in this study we use geometric morphometric techniques to objectively quantify leaf shape based on two leaf shape categories previously described by Shull and Iannetta (Fig. 1B), shape descriptors, climate factors, and climate regions (Fig. 1A) and investigate leaf shape across the United States over a 100-year period. We develop a shape analysis pipeline using pseudo-landmarks for this study, that uses leaf outlines (Fig. 1D) from *C. bursa-pastoris* herbarium specimens. We model how climate affects key leaf shape parameters at different temporal and spatial scales to thoroughly investigate the environmental factors shaping trait distribution.

**Figure 1.**
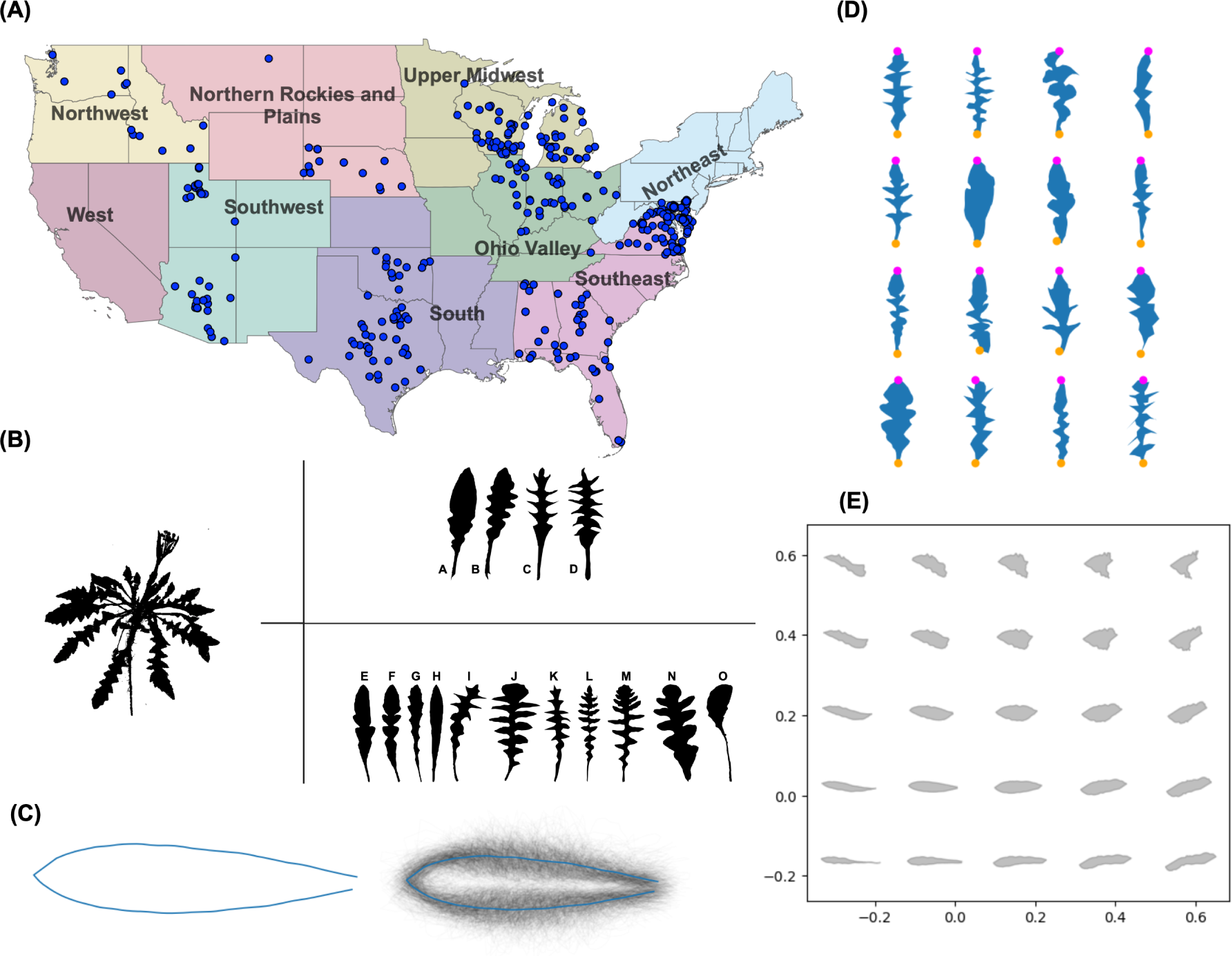
Overview of herbarium specimen selection, leaf shape types, and leaf shape analysis. (A). Map of the continental United States colored by climate region. Blue points represent herbarium specimen collection locations. (B). Schematic of leaf shape types. The left panel includes a representative of the *C bursa-pastoris* rosette taken from a herbarium specimen. [A-D]: Shull leaf shape types Simplex, Rhomboidea, Tenius, and Hetersis. [E-O]: Iannetta leaf shape types [E-H]: 1a-1d, [I-J]: 2b-2b, K: 3/4, L: 5, M:6, [N-O]: 7a-7b. (C). Mean leaf shape generated by Generalized Procrustes Analysis. The left leaf (blue outline) is the overall mean leaf shape and the right leaf is each individual leaf outline overlaid together in black with the mean leaf overlaid in blue. (D). Schematic of leaves included in leaf shape analysis, including true landmarks. Outlines of a representative sample of leaves (n = 12) included in this study are presented in blue. The two true landmarks, the leaf tip and leaf base, are represented by purple and orange points respectively. (E). Morphospace of theoretical leaves generated by inverse PCA. The morphospace projects five columns and rows of theoretical leaves generated by inverse PCA from leaf outlines included in this study.

## Materials and Methods

### Specimen collection and leaf outlines

We examined differences in leaf shape across the continental U.S and over a 100-year timespan (1921 - 2021) using 523 herbarium specimens of *C. bursa-pastoris* (Table S1). Each herbarium sample was accessed and downloaded from the Consortium for Midwest Herbaria online catalog (Midwest Herbaria, 2024). We only included samples with legible labels allowing us to identify the geographic location where each specimen was collected. To control for developmental differences in rosette development, only samples that were flowering when collected were included. Each state in the continental U.S. was assessed individually for sample availability and needed to have at least five potentially usable samples to be included in this study. Our final list of states includes Alabama, Arizona, Delaware, Florida, Georgia, Idaho, Illinois, Indiana, Maryland, Michigan, Montana, Nebraska, Nevada, Ohio, Oklahoma, Oregon, Texas, Utah, Virginia, Washington, and Wisconsin. All NOAA defined U.S. climate regions (Karl & Koss, 1984), except for the West, were represented in this study.

During the second selection step, each specimen was required to include one leaf separated from the whole plant and other leaves, with enough white space to easily outline that leaf. Our final data set included 497 leaves. A condensed list of specimens collected, including their climate regions can be found in Table S1 and an expanded list of all samples used in this study including the herbarium, label, and climate information can be found on GitHub (see Data Availability for information). Each leaf was outlined using the segmented line tool in ImageJ (Schindelin *et al*., 2012). Points each were included for both the right and left sides of each leaf, starting at either the right or left end of the petiole, around the leaf, and to the opposite end of the petiole. Each leaf was then saved as an XY coordinate text file. For each leaf, the area, perimeter, length (from tip to visible petiole base), and width were recorded using the ImageJ measurement tool with the settings area, shape descriptors, and perimeter selected.

### Data preparation and Generalized Procrustes Analysis

We analyzed each outlined leaf shape’s coordinate file with a shape analysis pipeline in Python using Jupyter notebook (Kluyver *et al*., 2016). This pipeline included importing leaf outline as coordinate text files, interpolating all points, and performing Generalized Procrustes Analysis (Procrustes distance). To perform landmark analysis, we first needed to orient each leaf so that each leaf was rotated and facing the same direction. To do this, we first found the indices (coordinate values/points) that represented the tip and base of each leaf. These indices were then re-indexed so that each leaf began at the base. Each leaf was rotated so that all leaf tips and leaf bases were facing the same direction. Due to the variability of *C. bursa-pastoris* leaf shape, we could only include two true landmarks for landmark analysis - the tip and the base of each leaf. Therefore, we assigned pseudo-landmarks from leaf tip to leaf base (left side of leaf) and then from leaf base to leaf tip (right side of leaf) so that each leaf included the same number of points. We then performed GPA on these re-indexed shapes. During GPA, each leaf was scaled and transformed to be compared to an arbitrary starting leaf (the first leaf in our dataset). After transformation, Procrustes distance is calculated and a mean leaf is generated. This process iterates across all leaves in our data set until a Procrustes threshold is reached.

The final products of GPA include a final Procrustes distance and a new set of Cartesian coordinates based on the scaled and transformed leaves. From GPA, we produced a mean leaf for the continental U.S. (Fig. 1C). We defined archetypal leaves representing the four Shull leaf shape types (Shull, 1909) and the seven Iannetta et.al. shape types (Iannetta *et al*., 2007). We then used GPA to match each leaf in our study to an archetypal leaf from both type categories. The final products of this pipeline were a series of CSV files that included “best matches” for each of the type categories, circularity values, and aspect ratio values.

### Principal component analysis and shape descriptors

After performing GPA, we performed principal component analysis (PCA) on the re-indexed leaves. We then performed inverse PCA to plot theoretical (eigen) leaves. Using the inverse PCA theoretical leaves, we defined a morphospace function to plot theoretical leaves from PC1 and PC2 eigenvalues along the PC space (Fig. 1E). We measured shape descriptors to describe differences in lobing and size between each leaf. We used circularity (circ), calculated as *circ* = (4*π × Area*) *÷ Perimeter* to measure lobing between leaves. In this equation, a value of 1 describes a perfect circle and values below 1 have increased lobes. We also used aspect ratio (ar) to measure changes in size (*ar* = *width ÷ length*) for each leaf. Lower aspect ratio values suggest a leaf is wider and shorter while higher aspect ratio values suggest a leaf is longer and narrower.

### Weather data collection

We collected average temperature (AT), maximum temperature (MAX), minimum temperature (MIN), and average precipitation (AP) for the location of each plant sample. We included three time-ranges in which we collected weather data:

**Date of collection (DOC)** = Climate on the date of collection

**Growing season (GS)** = Climate on the date of collection - Climate six months before DOC

**Year long (YL)** = Climate on the date of collection - Climate 365 days before DOC

To collect weather data, we generated a list of coordinates (latitude and longitude) for all specimens. We used the R package rnoaa (Edmund *et al*., 2014; Sparks *et al*., 2017) to download daily station data from the ghcnd database (Peterson *et al*., 1998). We then found up to 200 stations within a 50-mile radius of each location. We then separated out each set of stations by city and found all station ID information for each city. Using the filtered station IDs, we then found all TAVG, TMAX, TMIN, and PRCP data from 1920-01-01 to 2021-01-01 for each city. We used reported monthly TMAX and TMIN data to calculate YL TAVG. To find both the GS and AMB weather data points, we used the same process as above in addition to the R package zoo (Achim & Gabor, 2005) to find the beginning date of the previous six months or previous year.

### Statistical analysis

All statistical analyses were performed with R version 4.2.3 (RStudio Team, 2020; R Core Team, 2021). We used Pearson’s chi-square test of association to determine the strength of association between each leaf shape type category (Shull and Iannetta) and with climate region. We also modeled the interaction between shape descriptors using polynomial regression. We conducted one-sided t-tests and ANOVAs to determine associations between climate region and leaf shape.

Five polynomial regression models with h(degrees) of one to five were compared using standard parameters (k-fold cross validation of k=10). To estimate differences in leaf shape by shape descriptors, we performed an Analysis of Variance (ANOVA) on each climate x time model.

These models included:

**GS** = Shape Descriptor *∼ AT_GS_* + MAX*_GS_* + MIN*_GS_* + Climate Region

**YL** = Shape Descriptor *∼ AT_Y L_*+ MAX*_Y L_* + MIN*_Y L_* + Climate Region

**DOC** = Shape Descriptor *∼ AT_DOC_* + MAX*_DOC_* + MIN*_DOC_* + Climate Region

**IN***_GS_*= Shape Descriptor *∼ AT_GS_*\*AP*_GS_*

**IN***_Y L_* = Shape Descriptor *∼ AT_Y L_*\*AP*_Y L_*

**IN***_DOC_* = Shape Descriptor *∼ AT_DOC_*\*AP*_DOC_*

Where IN models included the interaction between the average temperature and average precipitation for each time-range. A parametric variance test, Tukey HSD, was performed to determine differences in shape descriptors between climate regions. We then performed Delta AIC model comparison (Mazerolle, 2023) to find the best model for explaining differences in variance between shape descriptors. We performed one-way ANOVA on shape descriptors to determine their respective associations with climate region using the following equations: *climate by circ* = *circ ∼ climate region* and *climate by ar* = *ar ∼ climate region*. We then performed a one-sided t-test on mean circularity and mean aspect ratio for each climate region.

## Results

### *Capsella bursa-pastoris* leaf shapes vary continuously

We first addressed whether *C. bursa-pastoris* leaves fall into distinct shapes, as previously found, or show continuous patterns of variation. We outlined 497 *C. bursa-pastoris* leaves collected from herbaria across the continental United States. We then analyzed each leaf outline (Fig. 1D) using a shape analysis pipeline generated for this study. Due to the high degree of intraspecies leaf shape variation, *C. bursa-pastoris* leaves so not show any discernible leaf margin architectural features in the mean leaf generated by Generalized Procrustes Analysis (Procrustes distance) (Fig. 1C). This lack of discernible features is also apparent in the eigen leaf (theoretical leaf) representations of the morphospace (Fig. 1E) as well. However, even though lobes are not represented in morphospace representations of the leaves, a pseudo-landmark approach still comprehensively measures the outline of the leaf (for example, see Chitwood *et al*. (2014a)). We used GPA to comprehensively compare each herbarium leaf outline to archetypal Shull (Fig. 1B[A-D]) or Iannetta types (Fig. 1B[E-O]), assigning leaves to categories based on the smallest Procrustes distance to an archetypal leaf. 94% (n = 470) of leaves best matched the “Rhomboidea” Shull type and 78% (n = 388) of leaves best matched the “Type 3” Iannetta type, consistent with previous common garden experiments that found that these were the most common leaf shape types (Shull, 1909; Iannetta *et al*., 2007; Hurka & Neuffer, 1997; Neuffer, 1990; Neuffer *et al*., 2018). Additionally, we measured leaf shape (lobing) using circularity (circ) and leaf size using aspect ratio (ar).

The morphospace PCA generated with the theoretical leaves from GPA but not the aspect ratio or circularity measurements revealed that leaf shapes vary continuously and there was considerable overlap in leaf shape (Fig. 2A,B). PC1 and PC2 explained 21% and 13% of the variance in shape respectively. Both the “Rhomboidea” and “Type 3” shape categories spanned a majority of the available PC space suggesting that focusing on shape types will miss a lot of within-type leaf shape variation (Fig. 2A,B). In addition, the “Rhomboidea” type encompassed the entire range of available shape descriptors (circularity and aspect ratio) values in this study (0.05758 to 0.76106 circularity values and 1.712 to 6.956 aspect ratio values) in addition to representing 94% of leaves in this study. Therefore, there is only one leaf shape type truly represented in this study, which prohibits between - shape type comparisons. Pearson’s chi-square test of association revealed that only the Shull leaf shape types were weakly correlated with climate region (Cramer’s V = 0.343, *p* = 2.03 *×* 10*^−^*^17^). This pattern of continuous variation, along with evidence that major shape types were found in every climate region and consistently across time, suggests that type is not the most effective way to investigate how the environment relates to leaf shape.

**Figure 2.**
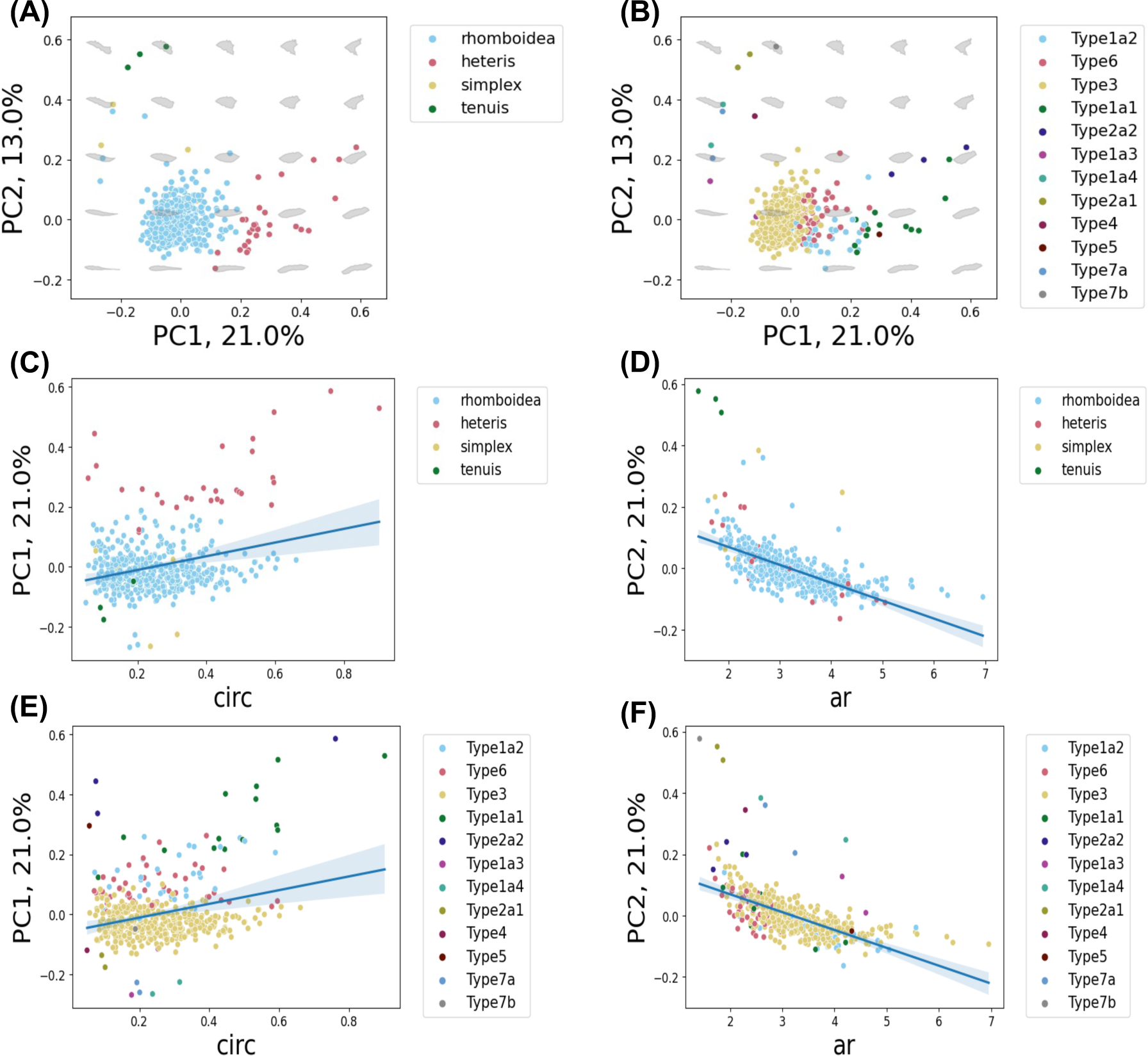
*C. bursa-pastoris* leaf morphospace, leaf shape types, circularity, and aspect ratio. (A). Morphospace PCA of leaves as classified by Shull leaf shape types. (B). Morphospace PCA of leaves as classified by Iannetta leaf shape types. (C,E). Graph of circularity (circ) against PC1. Leaves colored by their respective leaf shape type categories: Shull types (C) and Iannetta types (E). (D,F). Graph of aspect ratio (ar) by PC2, leaves colored by their respective leaf shape type categories: Shull types (D) and Iannetta types (F). The blue line represents the fitted linear regression and Tte gray band represents the 95% confidence interval.

The theoretical leaves of the morphospace PCA separate continuously along PC1 and are significantly associated with circularity (*p* = 9.14 *×* 10*^−^*^12^, Fig. 2C,E). The theoretical leaves also separate continuously along PC2 and are significantly associated with aspect ratio (*p* = 2 *×* 10*^−^*^16^, Fig. 2D,F)). Circularity and aspect ratio were also moderately positively correlated with each other(*Spearman^′^s ρ* = .302, *p* = 5.691 *×* 10*^−^*^1^.^2^). Polynomial regression showed a quadratic relationship between circularity and aspect ratio (*circ* = 0.00880 + 0.11880 *× ar −* 0.01287 *× ar*^2^, Fig 3A). There is strong constraint in change in circularity at extreme values of aspect ratio and more variation in circularity at intermediate values of aspect ratio. This pattern suggests that leaves can reach a maximum width (at low ar values) and a maximum length (at high ar values) only in highly lobed leaves, consistent with potential biological constraints for *C. bursa-pastoris* leaf shape.

**Figure 3.**
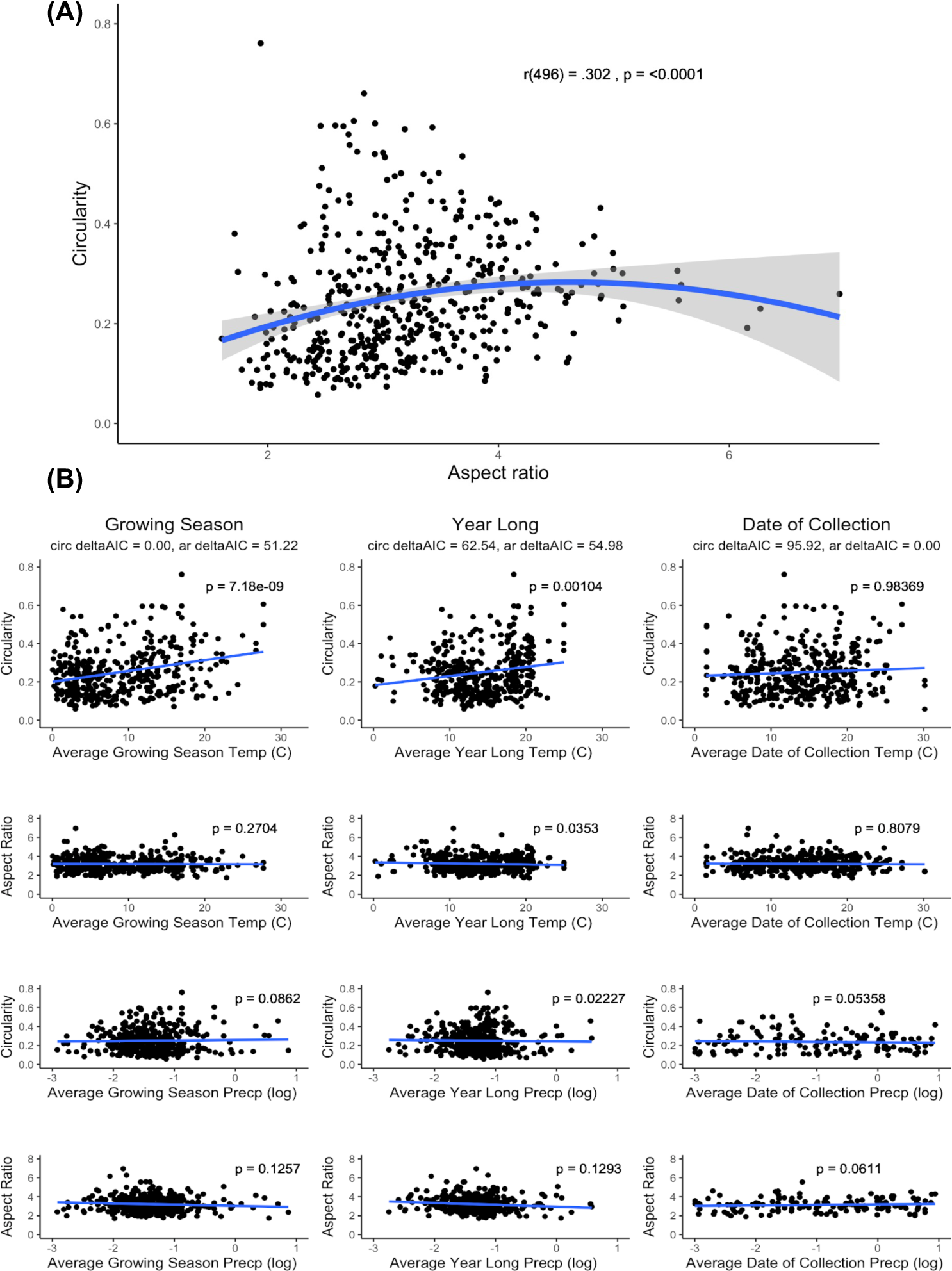
Modeling circularity and aspect ratio. (A). Circularity and aspect ratio exhibit a quadratic relationship. The blue line represents the fitted polynomial regression line. The gray band represents the 95% confidence interval. (B). Effects of climate on circularity and aspect ratio. The blue line represents the linear regression. The first column includes circularity and aspect ratio by the growing season (GS) climate conditions. The second column includes circularity and aspect ratio by the year long (YL) climate conditions. The third column includes circularity and aspect ratio by the date of collection (DOC) climate conditions. The model comparison deltaAIC is included for each climate x time model for both shape descriptors. The best model for explaining variance in circularity (lobing) was the GS model that includes climate region, with a deltaAIC score of 0. The best model for explaining variance in aspect ratio (size) was the DOC model including climate region.

Overall, the results of the geometric morphometric analysis suggest that both the Shull and Iannetta leaf shape types are less morphologically distinct than previously thought. Therefore, descriptive type categorizations are not meaningful for shape comparisons and will not be used going further in this study. Instead, we will focus on circularity and aspect ratio since they better describe the range of leaf shape variation on the PC and are correlated with climate region of origin.

### Leaf shapes vary by climate region and growing season temperature

To further investigate the relationship between leaf shape descriptors and climate region, we performed one-sided t-tests to determine if mean circularity and mean aspect ratio were individually significantly different between climate regions. The one-sided t-tests revealed significant differences among climate regions for mean circularity (*p* = 3.097 *×* 10*^−^*^08^) and mean aspect ratio (*p* = 2.294 *×* 10*^−^*^10^). We then performed one-way ANOVA and posthoc tests to determine which regions were significantly different from each other by circularity and aspect ratio (Fig. S2). Circularity was significantly different between the South and Northeast (*p* = 0.0000014), South and Southeast (*p* = 0.0000129), and South and Upper Midwest (*p* = 0.0076508). Aspect ratio was significantly different between the Upper Midwest and Northeast (*p* = 0.0044644). Overall, these result suggest that leaf shape differs broadly across the region, leading us to investigate the environmental factors that could contribute to this variation.

To test which environmental factors best explained phenotypic variation in leaf shape, we modeled shape descriptors as a function of average temperature (AT), maximum temperature (MAX), minimum temperature (MIN), and average precipitation (AP). Additionally, we investigated temperature at three time scales: the climate of the six months preceding collection (growing season, or GS), the climate of the year before collection (year long or YL), and climate on the date of collection (DOC). We compared the growing season and year-long models because previous work has shown that the environmental conditions of the specific time of year in which *C. bursa-pastoris* grows is more useful for determining the ecological niche than year-long data (Wilson Brown & Josephs, 2023). For this study, the DOC model acts as a negative control, as we do not expect the climate on the date of collection to affect leaf shape variation.

We used AIC model selection to determine which model best explained the variance in circularity and aspect ratio across the continental United States (Fig. 3B). The best fit model for explaining variance in circularity included every parameter in the GS model with no interaction effects. In this model, circularity increased as the average temperature (*p* = 7.15 *×* 10*^−^*^10^) and maximum temperature increased (*p* = 5.38 *×* 10*^−^*^12^). The second-best model was the YL model including every parameter with no interaction effects (p = 0.00153). The DOC and interaction models showed no significant differences in circularity across any of the included parameters. For aspect ratio, the DOC model was the best fit model and included every parameter. There were no significant associations between any of the temperature or precipitation variables and aspect ratio in the DOC model. There was a significant association between climate region and aspect ratio (p = 0.0120) in the DOC model.

### Growing season temperature explains leaf shape variation throughout the continental U.S and by region

Model selection revealed that the temperature in the six months before collection (GS) explains the variation in leaf shape better than the year long temperature (YL). However, the relationship between GS temperature and leaf shape is not consistent across the continental U.S. The South and Southeastern regions have the strongest associations between circularity (lobing) and average temperature (Fig 4A, S3). In the additional six climate regions, there was weak to no correlation between circularity and temperature. The largest range of circularity values was seen in the South (0.0951 to 0.7611) and and Southeast regions (0.0711 to 0.6057). The large range in circularity and strong association between temperature and shape could be due to a larger sample size in the Southeast but not the South. The Southeast included 152 individuals, the South - 78 individuals, the Upper Midwest - individuals, Ohio Valley - 57 individuals, Northeast - 51 individuals, Southwest - 40 individuals, Northern Rockies and Plains - 20 individuals, and the Northwest included 16 individuals. A summary of individuals by climate region is included in table S2.

**Figure 4.**
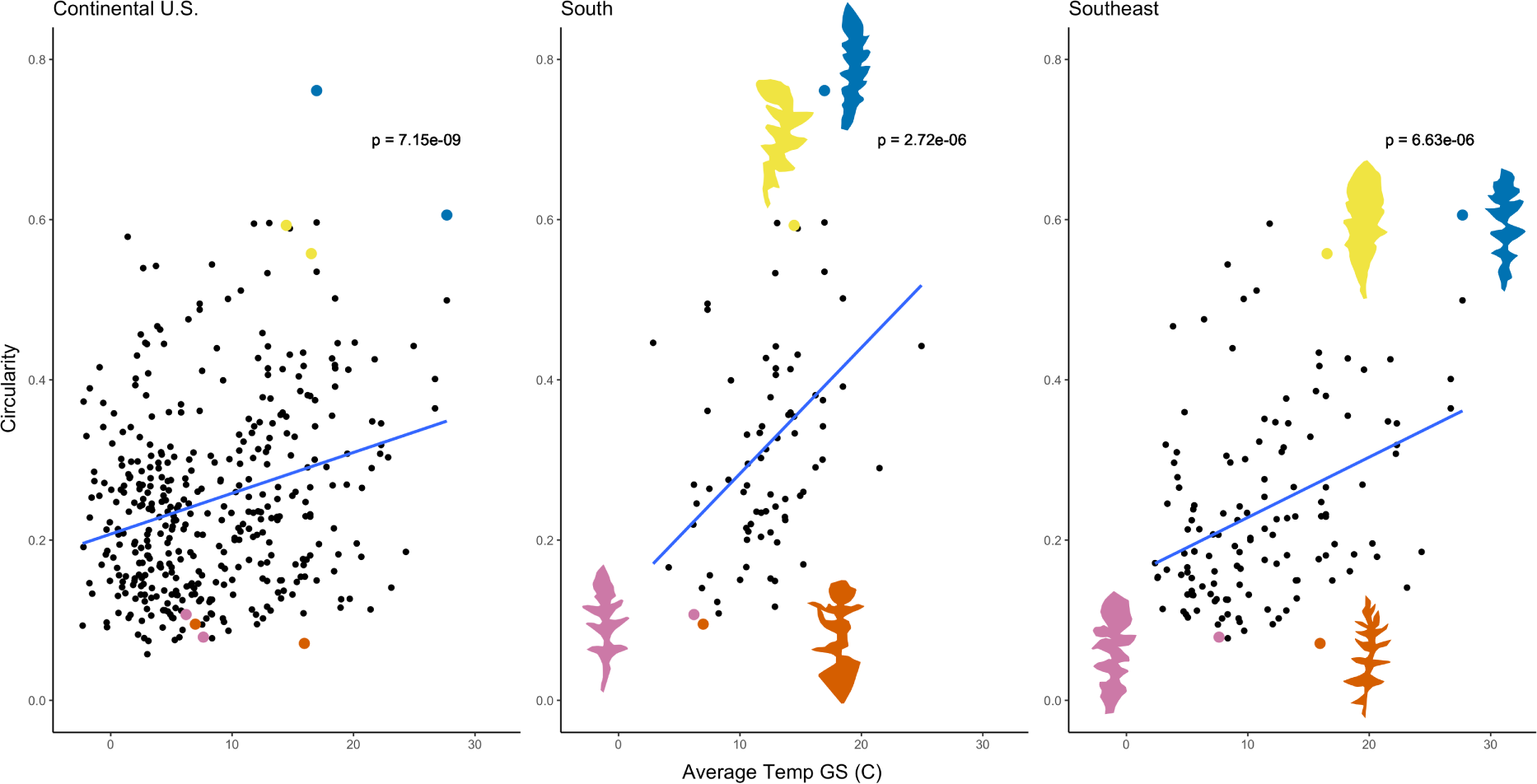
The relationship between average growing season (GS) temperature and circularity across all samples (left), in the South (middle) and in the Southeast (right). In all panels, the blue line represents the fitted linear regression. The two highest and two lowest circularity values for the south and southeast regions are colored in all three panels and represented by leaf images. Blue = highest circ, yellow = second highest circ, pink = second lowest circ, orange = lowest circ.

### Leaf shape variation has remained consistent over a 100-year time period

Leaf shape has not changed over time across the continental US although there were some changes within climate region. Circularity increased over time in the South (*p* = 1.08 *×* 10*^−^*^08^) and Southwest (p = 0.00683) regions while circularity decreased over time in the Northwest (p = 0.00628), Northern Rockies and Plains (p = 0.02929), Upper Midwest (p = 0.02093), and Southeast (p = 0.03362). Aspect ratio values followed a similar trend, where there was no change in aspect ratio over time across the continental U.S. and an increase in aspect ratio in the Upper Midwest (*p* = 6.69 *×* 10*^−^*^05^) and Northwest (p = 0.0225). Overall, leaf shape variation has been sustained over a 100-year period at the continental scale(Fig. S4).

## Discussion

In this study we found tremendous leaf shape variation within *C. bursa-pastoris* using tools that allowed us to systematically measure shape variation in scanned herbarium samples. We showed that this variation is not well-described by previous classification systems and, instead, propose that quantitative measures of lobing are the best way to quantify shape in this species. We linked this leaf shape variation to environmental variation and showed that this relationship, while significant across the North American range, is driven by associations within specific regions. While shape varied in space, we did not see significant changes in shape or the extent of variation in shape across time. Our results have clear implications for identifying the environmental factors contributing to intraspecies variation, as well as providing a guide for how one can systematically investigate shape variation in species with variable leaf shapes.

Historically, categories of leaf shape types have been used to subjectively categorize leaves (Shull, 1909; Iannetta *et al*., 2007; Shi *et al*., 2019; Schrader *et al*., 2021; Zhang *et al*., 2019). In the *C. bursa-pastoris studies* (Shull, 1909; Iannetta *et al*., 2007) there have been both an over representation of one leaf shape type and intermediate types that do not fit into one or more categories. Our Procrustes distance-based results suggest that there is substantial shape variation within categories. Within the Rhomboidea type alone, there is the full range of circularity found in this study. Therefore, distinctions made by category types may not be as meaningful as distinctions made by quantitative factors like circularity, where different shape types may be represented by one circularity value (Parins-Fukuchi, 2018; Felsenstein, 1973; Quinteros *et al*., 2006)). However, differences in leaf shape types may become more pronounced with the addition of more samples.

Instead of shape categories, this study used a pseudo-landmark approach to investigate leaf shape. Traditional landmark analysis of complex leaf shapes like those of *C. bursa-pastoris* can be difficult as there are inconsistencies in trait features like lobing depth, lobe/leaflet number, and lobe/leaflet size that make it challenging to assign landmarks. The use of pseudo-landmarks allow for comparisons between landmark points regardless of the above inconsistencies in shape (Dujardin *et al*., 2014; Lawing & Polly, 2010). These approaches will be broadly useful since *C. bursa-pastoris* is not the only plant species with inconsistent leaf shapes. For example, *Arabidopsis lyrata* which has varying leaf serrations (Vergeer & Kunin, 2011), and *Cardamine hirsuta* which has varying leaf shape and leaflet number (Canales *et al*., 2010).

While herbaria provide a remarkable source of plant traits and other data, there are some limitations to the conclusions that can be made from this data. The current 497 samples included in this study are biased in their collection times and locations. Most samples were collected within, and around more urban areas and the majority of repeated collection sites and collection times resulted from class projects at universities (Table S3). This bias has been well documented in herbarium studies (Moerman & Estabrook, 2006; Loiselle *et al*., 2008; Daru *et al*., 2018; Meineke & Daru, 2021; Panchen *et al*., 2019; Williams & Pearson, 2019) and highlights the need for repeated and sustained collections over an expanded collection range. In addition, traits measured from herbarium samples will be affected by both the genotype of the individual and the environment the individual grew in, making it difficult to distinguish what the underlying source of trait variation might be. Future work using common gardens, like that of Gupta *et al*. (2020), will be key for understanding how environment shapes leaf shape variation in *C. bursa-pastoris*.

As one of the most invasive plant species in the world, *C. bursa-pastoris* colonized, established, and flourished in a wide range of habitats and climates (Cornille *et al*., 2016, 2022; Wesse *et al*., 2021; Wilson Brown & Josephs, 2023). Some researchers have suggested that high plasticity may help *C. bursa-pastoris* persist across a wide range of environments (Choi *et al*., 2019; Cornille *et al*., 2022) For example, Choi *et al*. (2019) observed strong phenotypic plasticity for specific leaf area and leaf length in response to temperature and soil moisture in *C. bursa-pastoris*, and found evidence of selection for plasticity for specific leaf area. In addition, there is evidence that leaf type and traits like thickness and stomatal density vary genetically across the *C. bursa-pastoris* range (Neuffer *et al*., 2018). Here, we contribute to these previous results by showing that shape can be best described qualitatively, and that leaf circularity correlates with climate and differs between climate regions. While associations between leaf shape and climate suggest that shape is related to fitness in different types of environments, future work directly linking leaf shapes to fitness is needed to comprehensively understand the ecological importance of this trait during invasion.

Observations of variation in leaf shape also suggests there is a genetic mechanism underlying leaf shape response to the environment, although we do not measure this directly in this study. Previous research on the genetic basis for Shull leaf shape types suggests that there are two Mendelian loci with two alleles each that control the elongation of primary lobes (allele A) and the division of lobes (allele B) (Neuffer, 1990; Neuffer & Meyer-Walf, 1996). However, this study found continuous variation in leaf shape which would suggest the genetic mechanism of patterning leaf margins is not Mendelian or that it is strongly affected by environmental factors that varied across samples. Recent studies into the genetics of leaf lobing in *Cardamine hirsuta*, *Capsella grandiflora*, *Capsella rubella*, and other members of the lineage I Brassicaceae family has revealed the importance of *REDUCED COMPLEXITY 1 (RCO)* (Barkoulas *et al*., 2008; Blein *et al*., 2008; Sicard *et al*., 2014; Koenig & Weigel, 2015; Gan *et al*., 2016; Streubel *et al*., 2018; Runions *et al*., 2017; Gupta & Tsiantis, 2018). For the Capsella genus, the *RCO-A* gene induces the formation of lobes and reduces the blade surface area. In *C. grandiflora* specifically, *RCO-A* expression increases dramatically in low temperatures, almost ten times the normal expression at 20C (Sicard *et al*., 2014; Streubel *et al*., 2018). The *RCO-B* gene for both *C. grandiflora* and *C. rubella* induces the formation of serrations and is involved in the proximal - distal leaf patterning (Sicard *et al*., 2014; Streubel *et al*., 2018). *RCO* has yet to be characterized both genetically and functionally in *C. bursa-pastoris*. However, this work and other basic science studies are necessary first steps to understanding the biological mechanisms and potential consequences for both climate change and human intervention.

### Conclusion

In conclusion, our work has revealed that *C. bursa-pastoris* leaf shape exists on a spectrum and that discrete leaf shape types are more arbitrary than previously thought. We found that leaf shape is correlated with the growing season temperature of the plant, although this relationship varies among geographic regions. This suggests that climate has a large effect on leaf shape variation. Additionally, while our results do not show change in leaf shape over time, we do see the maintenance of leaf shape variation persist over the 100-year period included in this study. Finally, the use of herbarium samples and the leaf shape analysis pipeline created for this study has allowed us to compare complex, variable leaf shapes in an easy and less computationally intense way. This shape analysis pipeline will allow for further studies of complex shapes that were previously too difficult to pursue.

## Data availability

The data that support the findings of this study along with all code to do the analysis are openly available in a Github repository at the following link: https://github.com/AsiaH1994/Capsella Leaf Shape Herbarium project.

## Acknowledgments

We would like to thank Mila Vucelic for transcribing herbarium specimen labels and Miles Roberts for help with the weather data code.

## Author Contributions

ATH, DHC, and EBJ designed the research. ATH performed the research, data collection, and analysis and wrote the manuscript with advice from DHC and EBJ.

