## Supplemental Files for "Herbarium specimens reveal links between *Capsella bursa-pastoris* leaf shape and climate"

New Phytologist

### *New Phytologist* Supporting Information

The following Supporting Information is available for this article:

Files are in order as they appear in the text.

**Table S1 File: herb_paper_table_s1 .**

**Text: Table of Herbarium information for all *Capsella bursa-pastoris* specimens included in this study**. This includes the herbarium, collection number, barcode number, collector ID, climate region, state of collection, month, day, year, and Julian date of collection. All data was transcribed from available label data and associated data from the Consortium of Midwest Herbaria.

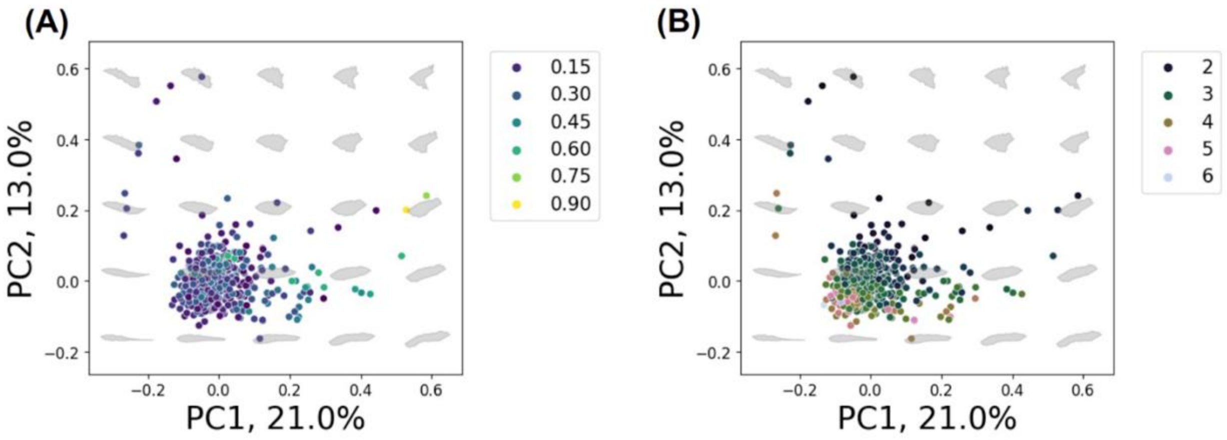


**Fig. S1 Circularity and aspect ratio cluster within the morphospace PCA**. A,B. Morphospace PCA of circularity (A) and aspect ratio (B). Circularity colors range continuously from blue (lowest circularity, most lobed) to yellow (most circular, least lobed). Aspect ratio colors range continuously from purple (lowest, wider and shorter shapes) to light blue (highest, thinnest and longest shapes).

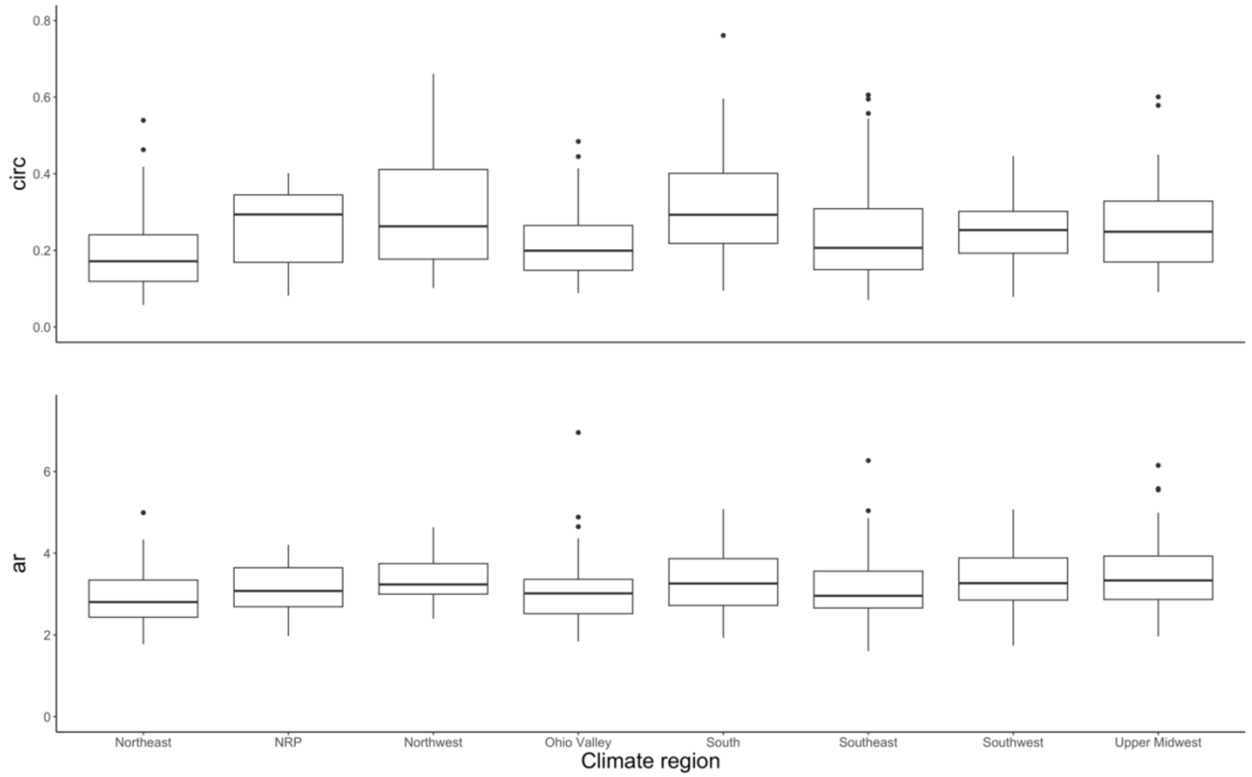


**Fig. S2 Circularity and aspect ratio values are significantly different between climate regions.** Top: boxplot of circularity values from the eight climate regions included in this study. Bottom: boxplot of aspect ratio values from the eight climate regions included in this study.

| Climate Region | Number of Individuals |
| --- | --- |
| Northeast | 51 |
| Northern Rockies and Plains | 20 |
| Northwest | 16 |
| Ohio Valley | 57 |
| South | 78 |
| Southeast | 152 |
| Southwest | 40 |
| Upper Midwest | 82 |

**Table S2** Sample sizes of individuals collected from each climate region.

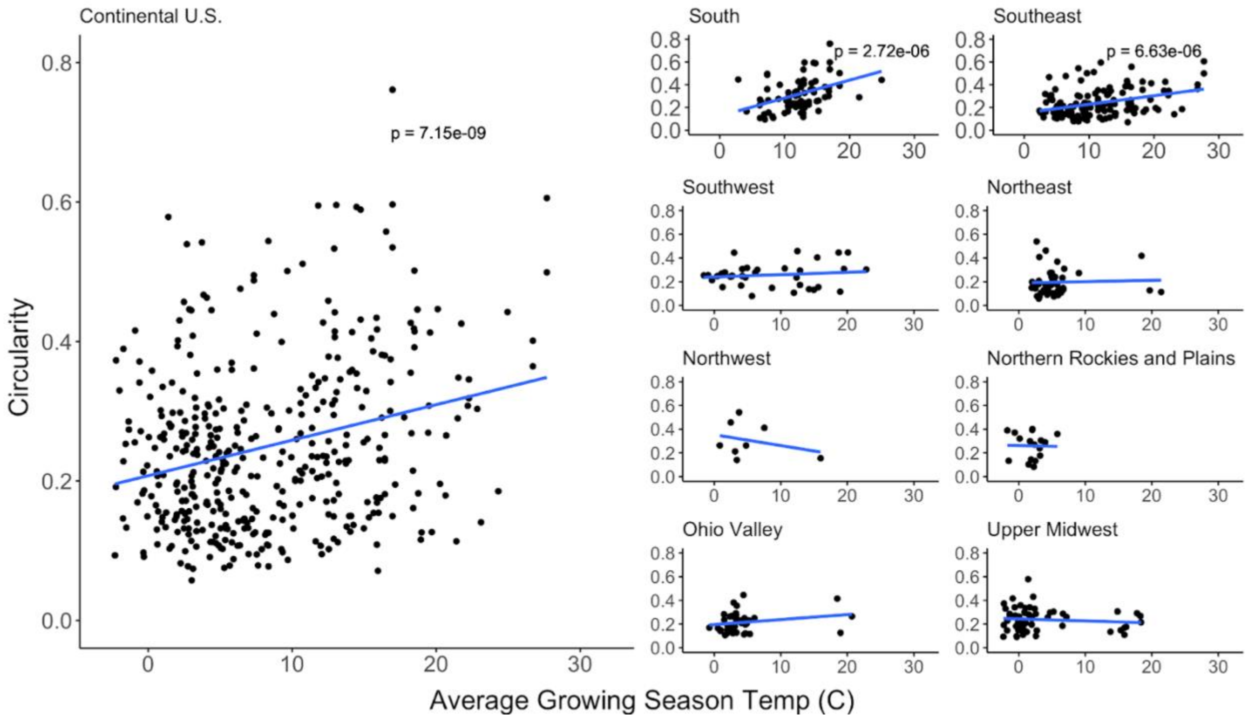


**Fig. S3 Circularity is strongly associated with average growing season temperature and by climate region.** Blue line represents fitted linear regression. P values for linear regression are provided for the continental U.S. and regions that are significantly associated with circularity over the average growing season temperature.

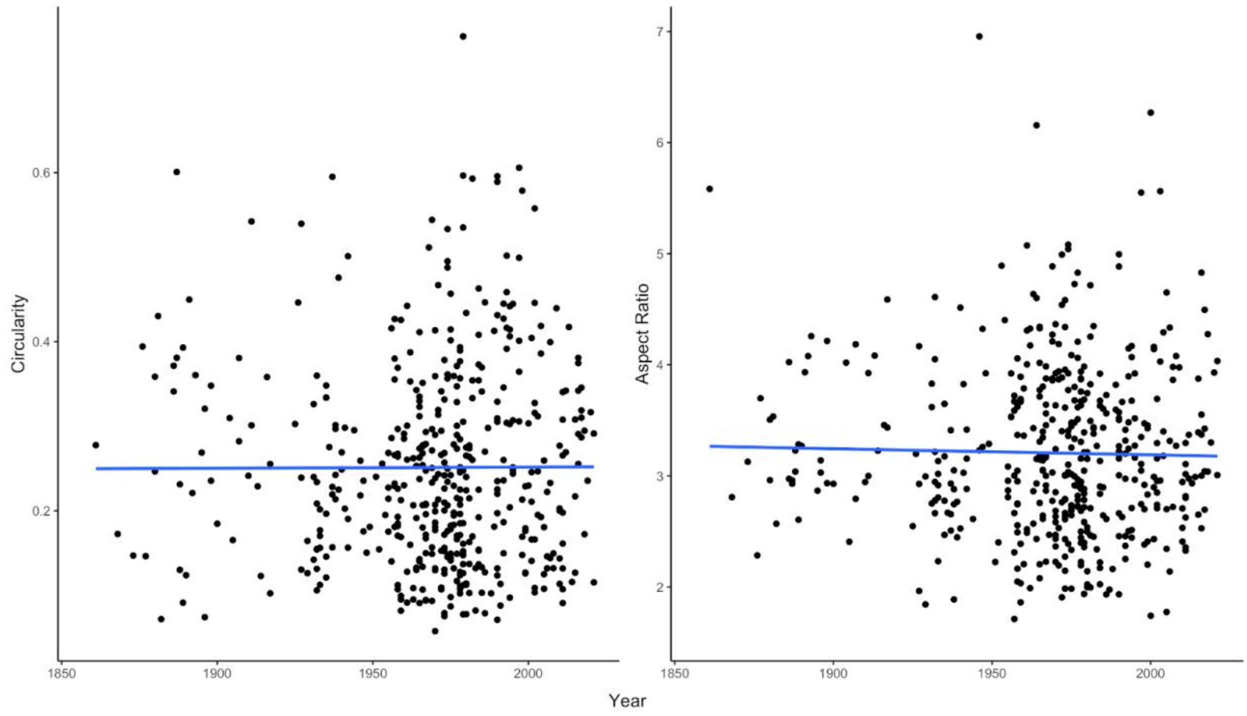


**Fig. S4 Circularity and aspect ratio have maintained variation over the 100-year period.** Circularity (left) and aspect ratio (right) graphed by year from 1890 – 2021. The blue line represents the fitted linear regression.

**Table S3** Individuals collected on or near a university campus.

**Table S3 File – herb_universities.xlsx**
